# Human Papilloma Virus does not fully inactivate p53 cellular activity in HNSCC

**DOI:** 10.1101/2025.02.11.637252

**Authors:** Jovanka Gencel-Augusto, Hua Li, Liam C. Woerner, Ashir A. Borah, Jeffrey N. Myers, Patrick Ha, Daniel E. Johnson, Jennifer R. Grandis

**Affiliations:** Department of Otolaryngology-Head and Neck Surgery, University of California San Francisco (UCSF), San Francisco, CA, USA; UCSF Helen Diller Family Comprehensive Cancer Center, San Francisco, CA, USA; Department of Head and Neck Surgery, MD Anderson Cancer Center, Houston, TX, USA

## Abstract

Head and neck squamous cell carcinoma (HNSCC) is a major global health challenge. Inactivation of the tumor suppressor p53 is the most frequent molecular event in this malignancy. p53 inactivation occurs either through *TP53* mutations in human papilloma virus (HPV)-negative cases or via HPV-mediated p53 degradation in HPV-positive (HPV+) cases, where most tumors retain a wild-type (WT) *TP53* allele. This underscores the critical role of p53-regulated processes in HNSCC pathogenesis. Clinically, HPV+ HNSCC is associated with significantly better outcomes than HPV-negative cases. However, despite HPV E6-mediated degradation of p53, approximately 10% of HPV+ HNSCC tumors harbor *TP53* mutations, suggesting an additional selective pressure to suppress p53 signaling.

In this study, we demonstrate that HPV+ *TP53*-WT HNSCC cells have residual, tumor suppressive p53 activity. Specifically, analysis of human tumor data reveals that among HPV+ HNSCC cases, those with WT *TP53* have significantly better survival outcomes than both HPV+ cases with *TP53* mutations and HPV-negative cases. Experimentally, genetic ablation of WT p53 in HPV+ HNSCC cells increased proliferation, migration, and invasion. Transcriptomic analysis revealed that p53 continues to regulate gene expression despite the presence of HPV.

Further, human tumors with HPV+ *TP53*-WT status exhibit tumor-suppressive methylation patterns, fewer chromosomal alterations, and suppression of PI3K-AKT signaling compared to HPV+ *TP53*-mutant tumors. Importantly, loss of WT p53 in HPV+ HNSCC cells increased the levels of PI3K catalytic subunit p110α, reduced expression of the molecular PI3K-AKT inhibitor INPP5D and enhanced sensitivity to pharmacologic PI3K inhibition.

Together, our findings challenge the prevailing view that p53 is completely inactivated in HPV+ HNSCC and reveal tumor suppressive, p53-driven mechanisms that persist in these tumors. These insights highlight a potential role for *TP53*-based stratification in guiding treatment decisions and suggest new therapeutic vulnerabilities in HPV+ HNSCC.

## Introduction

Head and neck squamous cell carcinoma (HNSCC) represents a significant global health challenge, with approximately 900,000 new cases diagnosed annually [1]. Nearly all cases involve inactivation of p53, a key regulator of anti-tumor responses. In ∼70% of HNSCC cases, p53 inactivation occurs through genetic alterations in *TP53*, while ∼20% result from human papillomavirus (HPV) infection (HPV-positive, HPV+), which promotes constitutive degradation of the p53 protein [2]. This underscores the central role of p53-regulated pathways in HNSCC pathogenesis. Clinically, *TP53* mutations in HNSCC are associated with poor response to conventional therapies and significantly worse survival outcomes [3]. Despite extensive efforts, no effective treatments exist for the vast majority of *TP53*-mutant cancers, largely due to the challenge of selectively targeting mutant p53 proteins, which are often deemed “undruggable”. In contrast, HPV+ HNSCC, in which 90% of cases retain wild-type (WT) *TP53* alleles but experience HPV-mediated p53 degradation, exhibits markedly better prognosis [2, 4] and therapeutic responsiveness than HPV-negative (mostly *TP53*-mutant) HNSCC [2]. The molecular determinants for these differences are not well understood.

p53 functions as a transcription factor, orchestrating the expression of hundreds of genes (both up and down) involved in apoptosis, cell cycle arrest, senescence, and several other processes, to suppress tumor development [5]. HPV promotes p53 degradation through the viral E6 protein, which hijacks the host E3 ubiquitin ligase E6AP, targeting p53 for proteasomal degradation [6]. This mechanism mirrors the cellular intrinsic regulation of p53 via MDM2, another E3 ubiquitin ligase that tags p53 for degradation [7]. Notably, p53 transcriptionally activates *MDM2*, establishing a tightly regulated feedback loop that restrains excessive p53 activity [7], which could otherwise be detrimental to normal cells.

Low p53 protein levels are important for cellular and organismal homeostasis. Seminal studies have demonstrated that MDM2-mediated regulation of p53 signaling is essential for maintaining physiological health. Genetic deletion of *Mdm2* in mice results in p53-dependent embryonic lethality [8]. Similarly, conditional deletion of *Mdm2* in adult murine tissues triggers robust activation of p53 target genes that result in severe multi-organ toxicities - phenotypes that are rescued by *p53* deletion [9]. This evidence underscores the key role of MDM2 in regulating p53 activity to prevent irreversible cellular outcomes. Consequently, under homeostatic conditions, healthy tissues maintain very low p53 protein levels due to MDM2-driven degradation. However, low p53 protein abundance does not equate to a complete absence of p53 signaling [10]. Beyond the well-characterized roles of p53 in acute responses to cellular stressors such as DNA damage or oncogene activation (i.e. apoptosis, cell cycle arrest), p53 also performs essential homeostatic functions. These functions include preserving mesenchymal stem cell integrity [11], regulating airway epithelial differentiation [12, 13], and modulating cellular metabolism [14, 15], among other fundamental processes that contribute to organismal function and tumor suppression. Taking these data together, we hypothesized that HPV+ *TP53*-WT HNSCCs that maintain low p53 protein levels due to HPV presence retain residual p53 tumor suppressive signaling.

To investigate this, we first stratified human HPV+ HNSCC cases by *TP53* status, revealing that HPV+ *TP53*-WT cases have significantly better survival outcomes than HPV+ *TP53*-mutant and HPV-negative cases. Using a panel of HPV+ *TP53*-WT HNSCC cell lines with or without p53 ablation, we performed downstream assays. Interestingly, further suppression of p53 through RNA (RNAi) or CRISPR interference (CRISPRi) in HPV+ HNSCC cells promoted more aggressive cellular behaviors, suggesting the presence of residual tumor-suppressive p53 activity that may constrain tumor progression despite HPV infection. RNA sequencing revealed p53-driven transcriptional regulation of multiple genes both up and down. Several of these genes were also differentially expressed upon ectopic WT p53 expression in HPV-negative HNSCC cells, reinforcing that the observed transcriptional programs in HPV+ HNSCC are indeed p53-dependent. Human HPV+ *TP53*-WT HNSCC tumors exhibit increased gene methylation in oncogenic pathways, fewer chromosomal alterations, and lower PI3K-AKT signaling activity compared to HPV+ *TP53*-mutant tumors, further supporting the persistence of p53 signaling in the presence of HPV. Finally, we reveal that p53 functions as an endogenous suppressor of the PI3K-AKT pathway and that its loss renders HPV+ HNSCC cells more sensitive to pharmacologic PI3K inhibition.

Our study provides evidence of a residual tumor-suppressive program driven by p53 in HPV+ HNSCC, challenging the prevailing notion that p53 function is completely inactivated in HPV-infected cells. Our findings suggest that this residual activity may contribute to the differential clinical outcomes between HPV+ and HPV-HNSCC. We propose that stratification of HPV+ HNSCC cases by *TP53* status could improve responses to current therapies. Furthermore, we propose that HPV+ HNSCC models serve as a unique platform to study disease-specific p53-driven tumor suppression mechanisms that may inform novel therapeutic strategies for *TP53-*mutant cancers.

## Results

To establish the clinical and molecular context for our study, we first analyzed *TP53* status and patient outcomes in head and neck squamous cell carcinoma (HNSCC) using The Cancer Genome Atlas (TCGA, PanCancer Atlas dataset). Consistent with prior reports [16], HPV+ HNSCC cases predominantly retain WT *TP53*, while HPV-negative cases are frequently characterized by *TP53* mutations (Figure 1A). Moreover, patients with HPV+ HNSCC exhibit significantly improved overall survival compared to those with HPV-negative disease (Figure 1B). Notably, the mechanism by which HPV promotes p53 degradation via the E6/E6AP complex closely mirrors the cellular regulation of p53 through MDM2, another E3 ubiquitin ligase that targets p53 for proteasomal degradation (Figure 1C). We hypothesized that despite viral-mediated p53 degradation, residual p53 activity in HPV+ tumors may contribute to a more favorable prognosis.

**Figure 1.**
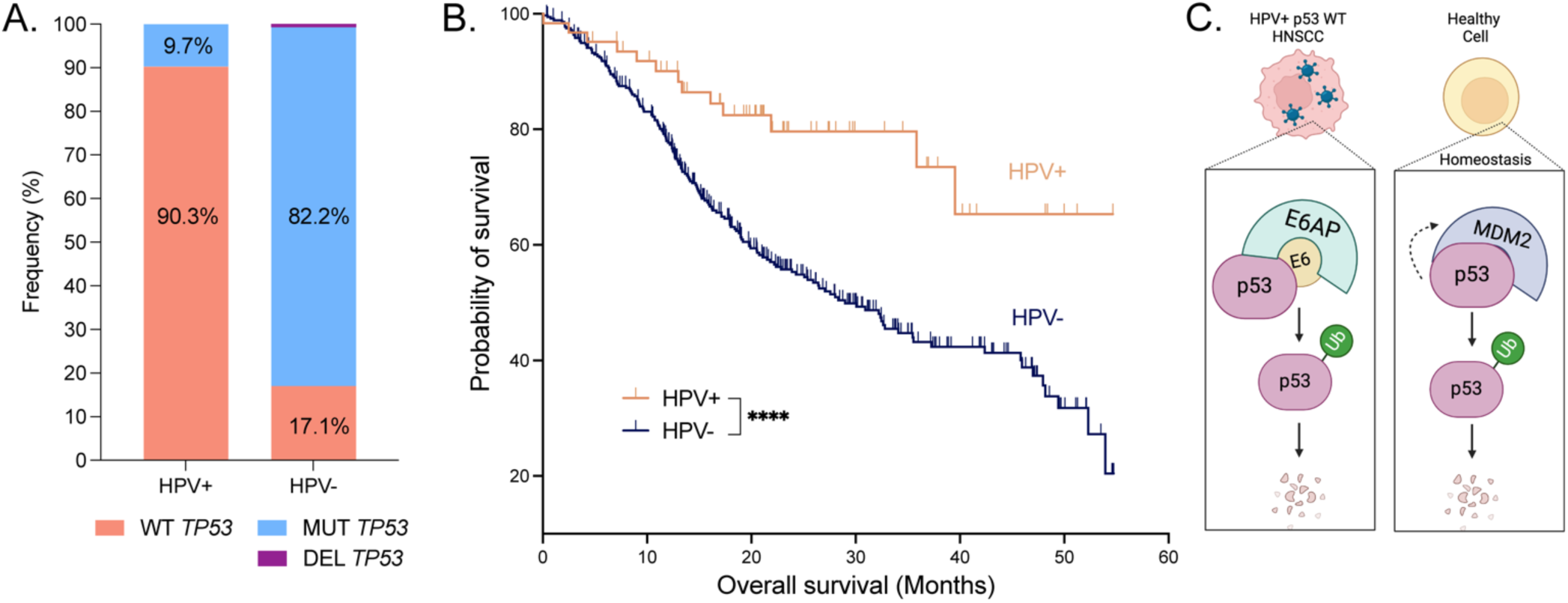
HPV+ HNSCC: Survival, *TP53* status, and parallels to physiological p53 regulation. **A.** Frequency of p53 (*TP53*) alterations in HNSCC. MUT=mutant, DEL=deleted. **B.** Overall survival comparison between HPV-positive (HPV+) and HPV-negative (HPV-) HNSCC. Data in A and B obtained from The Cancer Genome Atlas (TCGA, cBioPortal). ****p<0.00005. **C.** Diagram illustrating the molecular parallels between HPV-mediated p53 degradation and MDM2-mediated p53 degradation.

### Retention of WT p53 confers even better outcomes to HPV+ cases compared to HPV-negative HNSCC

We first stratified HPV+ HNSCC tumors from TCGA by *TP53* status (WT vs. mutant) and assessed clinical outcomes. Interestingly, HPV+ *TP53*-WT tumors exhibited significantly improved progression-free survival compared to both HPV+ *TP53*-mutant cases and HPV-negative cases (Figure 2A). Notably, survival outcomes in HPV+ *TP53*-mutant cases were not significantly different from those in HPV-negative cases.

**Figure 2.**
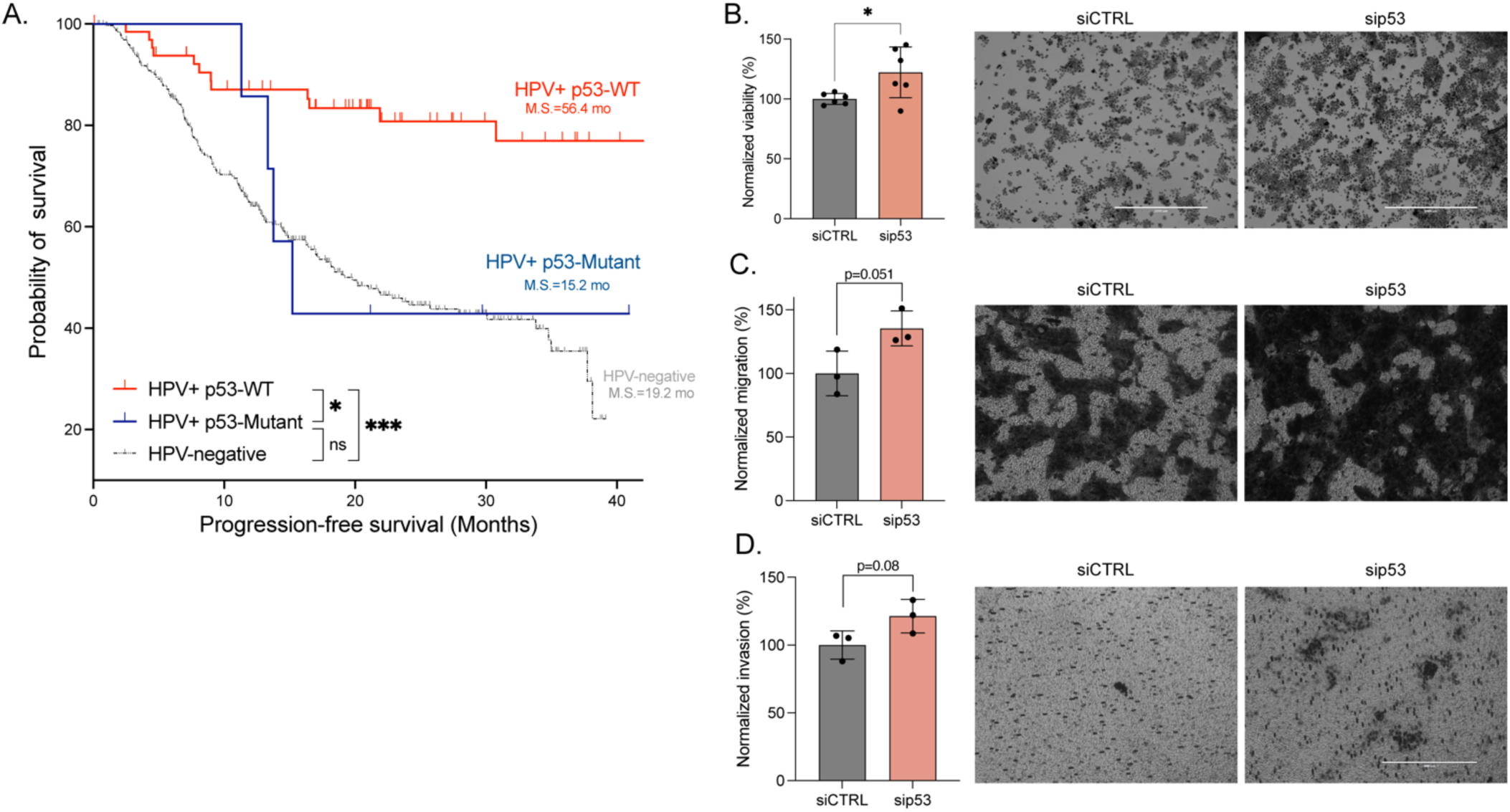
Retention of WT *TP53* is associated with improved survival and suppressed tumorigenic phenotypes in HPV+ HNSCC. **A.** Progression-free survival analysis comparing HPV+ *TP53*-WT (N=64), HPV+ *TP53*-Mutant (N=7), and HPV-negative (N=415) HNSCC cases. *p < 0.05, ***p<0.0005, ns=not significant. MS=median survival in months (mo). **B.** Viability assay following *p53* knockdown (48h, UM-SCC-47), *p<0.05. Migration **(C)** and invasion **(D)** assays in UM-SCC-47 following *p53* knockdown (48 hours). Representative end point images are presented to the right of the corresponding bar plots.

Based on these findings, we investigated the functional consequences of p53 ablation in *TP53*-WT HPV+ HNSCC cell lines using siRNA-mediated p53 knockdown, with a non-targeting siRNA as a control. To assess the phenotypic impact of p53 loss, we examined cell proliferation, migration, and invasion. Treatment with p53 siRNA effectively reduced p53 mRNA and protein levels compared to a non-targeting control siRNA at the assessed time points (Supplementary Figures 1A and 3). p53 knockdown significantly enhanced proliferation in the HPV+ human HNSCC cell lines UM-SCC-47 (Figure 2B), UPCI-SCC-090, and UD-SCC-2 (Supplementary Figure 1B). We also used UM-SCC-47 cells harboring dCas9 (UM-SCC-47-dCas9) to assess the impact of p53 knockdown on cell proliferation through CRISPR-interference. This approach also showed increased cell proliferation upon p53 ablation (Supplementary Figure 1C-D).

Additionally, in UM-SCC-47, p53 depletion increased both migration and invasion capabilities (Figure 2C-D). UPCI-SCC-090 and UD-SCC-2 did not exhibit baseline migration or invasion, preventing assessment of these processes upon p53 knockdown.

These findings suggest that WT p53 activity present in HPV+ HNSCC cells mitigates tumor cell “progression” (proliferation, migration, invasion), highlighting a tumor-suppressive role despite HPV-mediated p53 degradation.

### Functional p53 transcriptional programs persist in HPV+ HNSCC

Given the phenotypic consequences of p53 ablation in HPV+ HNSCC cells, we hypothesized that p53 transcriptional activity is present. To investigate this, we performed RNA-sequencing following p53 knockdown in two HPV+ *TP53*-WT HNSCC cell lines, UM-SCC-47 and UPCI-SCC-090, using a non-targeting siRNA as a control. Samples were collected 30 hours post-transfection, a time point selected based on the kinetics of *CDKN1A* (p21, a canonical p53 target gene) expression changes observed at 8, 12, 24, 30, and 48 hours post-transfection (Supplementary Figure 2). *CDKN1A* levels were significantly reduced upon p53 knockdown across all time points, with the most pronounced effect at 30 and 48 hours. We selected 30 hours to minimize the influence of compensatory signaling that is not p53-dependent.

Differential gene expression analysis revealed many differentially expressed genes (DEGs; p<0.01; |Log_2_fold-change| > 0) following p53 knockdown in both cell lines (Figure 3A-B, Supplemental Table 1). Among the identified DEGs, ∼60% were downregulated and ∼40% were upregulated. These findings are consistent with previous reports of p53 as both transcriptional activator and repressor [17]. To determine whether these DEGs include direct p53 targets, we compared our DEG lists with previously identified p53-bound genes from p53 chromatin immunoprecipitation sequencing (ChIP-seq) datasets [18]. Approximately 20% of the DEGs in each cell line overlapped with known direct p53 targets (Figure 3C, Supplemental Table 1). However, the total number of DEGs was markedly lower than what has been reported in other cell types without HPV infection, either following *Mdm2* genetic deletion [19] or in response to DNA damage-induced p53 activation [18]. This suggests that while p53 functionality is restricted in HPV+ HNSCC, it remains present.

**Figure 3.**
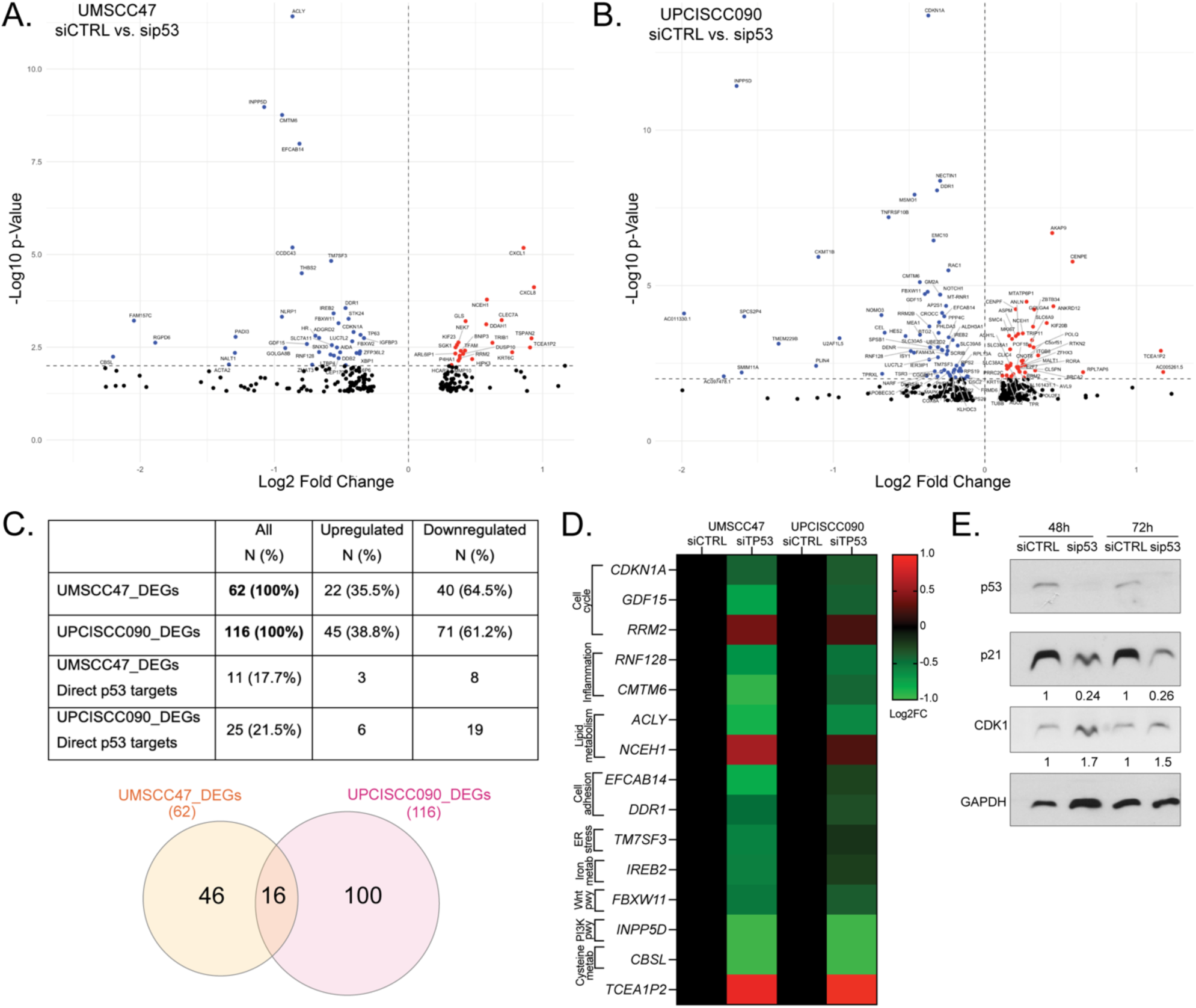
p53 regulates gene expression in the presence of HPV. **A.** Volcano plot shows significant differentially expressed genes (DEGs, blue=downregulated, red=upregulated, p<0.01) in UM-SCC-47 upon *p53* knockdown (two biological replicates per experimental condition). **B.** Volcano plot for DEGs in UPCI-SCC-090 upon *p53* knockdown. **C. Top:** Table shows a summary of total, upregulated, or downregulated DEGs per cell line, including number of genes that are previously reported as direct p53 targets (“p53-bound”) [18]. **Bottom:** Venn diagram shows shared and cell line-specific DEGs in UM-SCC-47 and UPCI-SCC-090. **D.** Heatmap illustrates average expression levels of shared DEGs in both cell lines. Green shades indicate downregulated genes, while red shades indicate upregulated genes. Values are represented as Log2 fold-change (Log2FC), with those exceeding |1| capped at +1 or −1 for illustrative clarity. Rows are genes and columns are experimental conditions. The cellular processes related to each gene are shown on the left. Pwy=pathway, Metab=metabolism. *TCEA1P2* is a pseudo-gene with no known function. **E.** Validation of protein changes, including p53 knockdown, p21 (*CDKN1A*) downregulation, and CDK1/2 upregulation in UM-SCC-47 at 48 and 72 hours post-transfection. Quantification of blots is shown relative to GAPDH and to siCTRL samples per time point, below each blot.

We further analyzed DEGs shared between the two HPV+ HNSCC cell lines and identified 16 common genes (Figure 3C, Supplemental Table 1 - red font), of which 13 were downregulated upon p53 knockdown, indicating p53-dependent activation (Figure 3D). Among these downregulated genes were *CDKN1A* (p21), *INPP5D*, and *GDF15,* which are established tumor-suppressors [20, 21]. Conversely, two of the three shared upregulated genes upon p53 knockdown, *RRM2* and *NCEH1*, have been previously described as tumor-promoting [22, 23]. Collectively, these 16 shared DEGs regulate diverse cellular processes, including cell cycle progression, inflammation, lipid metabolism, cell motility, endoplasmic reticulum (ER) stress, iron metabolism, and cysteine metabolism (Figure 3D).

Consistent with the mRNA-based findings, we confirmed that p21 protein was downregulated following p53 knockdown, persisting at 48 and 72 hours post-transfection. Similarly, CDK1/2, key regulators of cell cycle progression, were upregulated at both time points upon p53 ablation (Figure 3E).

Taken together, these findings demonstrate that WT p53 maintains a residual transcriptional program in HPV+ HNSCC, regulating genes involved in tumor suppression and cellular homeostasis despite HPV-mediated degradation.

### p53 transcriptional regulation in HPV+ HNSCC is recapitulated in HPV-negative HNSCC

To validate that the observed transcriptional changes in HPV+ cells result from p53-driven regulation, we first restored p53 protein levels by knocking down E6 (Supplemental Figure 3A). This approach effectively rescued the expression of p53 targets *CDKN1A* and *INPP5D* (Figure 4A; Supplemental Figure 3B).

**Figure 4.**
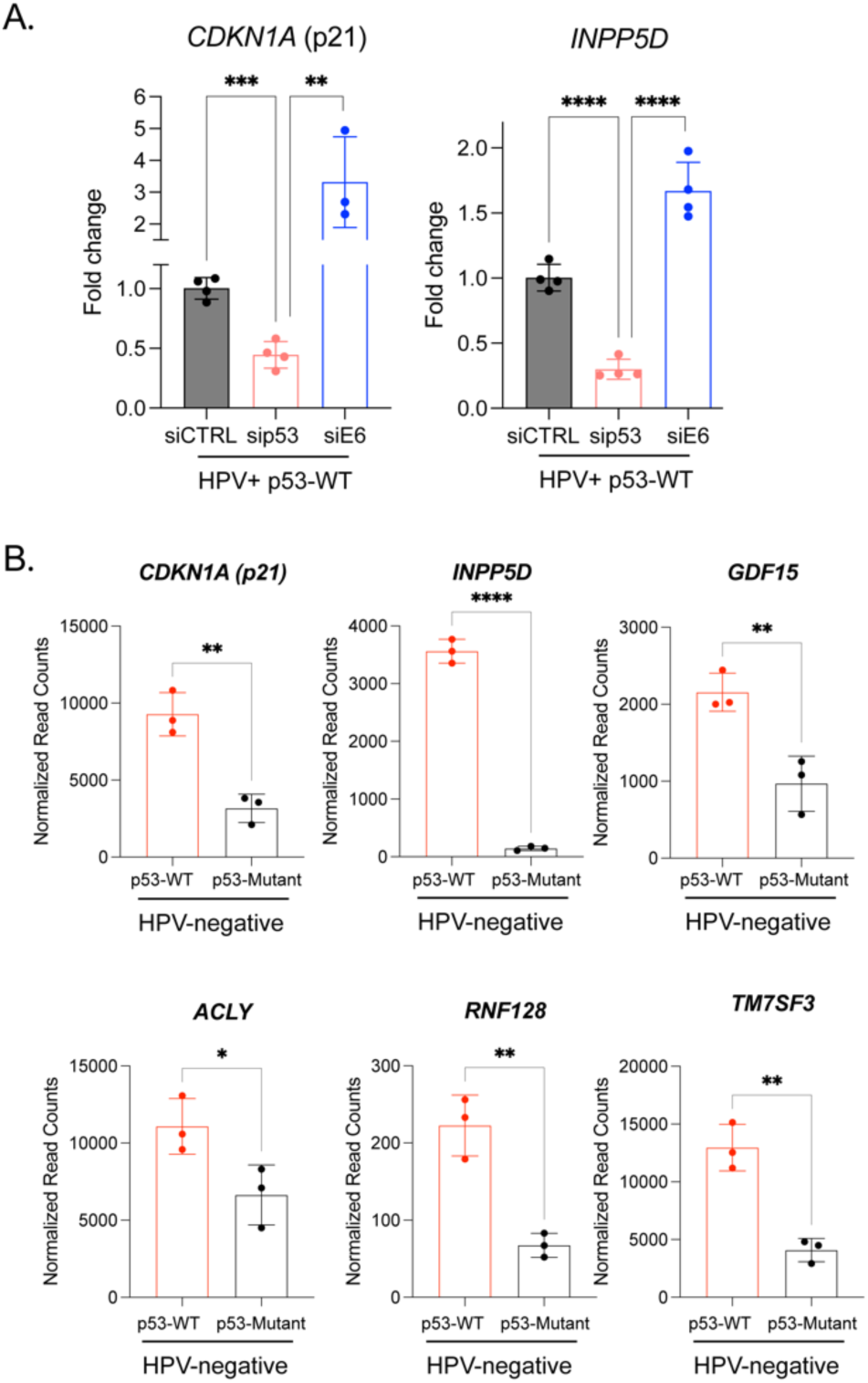
p53 transcriptional regulation in HPV+ HNSCC is recapitulated in the absence of HPV E6. **A.** mRNA expression levels of p53 target genes *CDKN1A* and *INPP5D* decrease upon p53 knockdown and are restored (and elevated) following E6 knockdown in UM-SCC-47 HPV+ HNSCC cells. **B.** Normalized RNA-sequencing read counts for *CDKN1A, INPP5D, GDF15, ACLY, RNF128, TM7SF3* in isogenic HPV-negative PCI-13 HNSCC cell lines with either p53-WT or p53G245D (loss-of-function mutant) expression.

To further confirm that these transcriptional effects are p53-dependent and not exclusive to HPV+ cells, we examined the expression of these genes in an HPV-negative context. We utilized PCI-13 HNSCC cell lines (harboring a spontaneous *TP53* deletion) engineered to stably express ectopic *TP53*-WT or *TP53*-mutant (p53G245D, loss-of-function mutant) [24]. Normalized read counts from RNA-sequencing analysis in these cell lines showed significantly higher expression levels of *CDKN1A*, *INPP5D*, *GDF15*, *ACLY*, *RNF128*, and *TM7SF3* (first shown in Figure 3D) in HPV-negative PCI-13 cells expressing WT p53 compared to their p53-mutant counterparts (Figure 4B).

These findings indicate that the transcriptional changes observed in HPV+ HNSCC cells (Figure 3) are specifically driven by p53.

### Significant molecular differences between HPV+ TP53 WT and HPV+ TP53-mutant human HNSCC

Based on our experimental findings suggesting tumor-suppressive signaling in HPV+ HNSCC cells that retain WT *TP53* alleles, we analyzed human tumor data. We stratified HPV+ HNSCC tumors from TCGA by *TP53* status (WT vs. mutant) (first presented in Figure 2A) and assessed molecular features. Despite the low prevalence (10%) of *TP53* mutations in HPV+ HNSCC, we identified several statistically significant molecular differences between *TP53*-WT and *TP53*-mutant HPV+ tumors.

We analyzed global RNA expression differences between *TP53*-WT and *TP53*-Mutant HPV+ HNSCC tumors (Supplementary Figure 4). Pathway enrichment analysis revealed that *TP53*-mutant tumors exhibit upregulation of genes associated with cell growth processes, including epithelial development, tissue morphogenesis, WNT signaling, and anabolic metabolism. In contrast, *TP53*-WT tumors showed increased expression of genes involved in immune regulation, DNA repair, and catabolic metabolism, among other processes. These data further support the tumor-suppressive role of p53 in HPV+ HNSCC.

In addition to gene expression differences, we also observed differential DNA methylation patterns between HPV+ *TP53*-WT and HPV+ *TP53*-mutant tumors. These data were obtained from the HM27 and HM450 arrays, DNA methylation profiling platforms available in cBioPortal for assessing epigenetic alterations in cancer [2]. These arrays cover CpG methylation in promoter, enhancer, and gene body regions, and identify methylation-driven gene silencing. We found that *TP53*-WT tumors exhibited significantly higher overall gene methylation compared to *TP53*-mutant tumors. Specifically, *TP53*-mutant tumors showed significantly increased methylation in only 23 genes, whereas *TP53*-WT tumors displayed elevated methylation across more than 500 genes (Figure 5A). Pathway enrichment analysis revealed that methylated genes in *TP53*-mutant tumors were predominantly associated with tumor-suppressive pathways, while methylated genes in *TP53*-WT tumors were linked to tumor-promoting pathways (Figure 5A).

**Figure 5.**
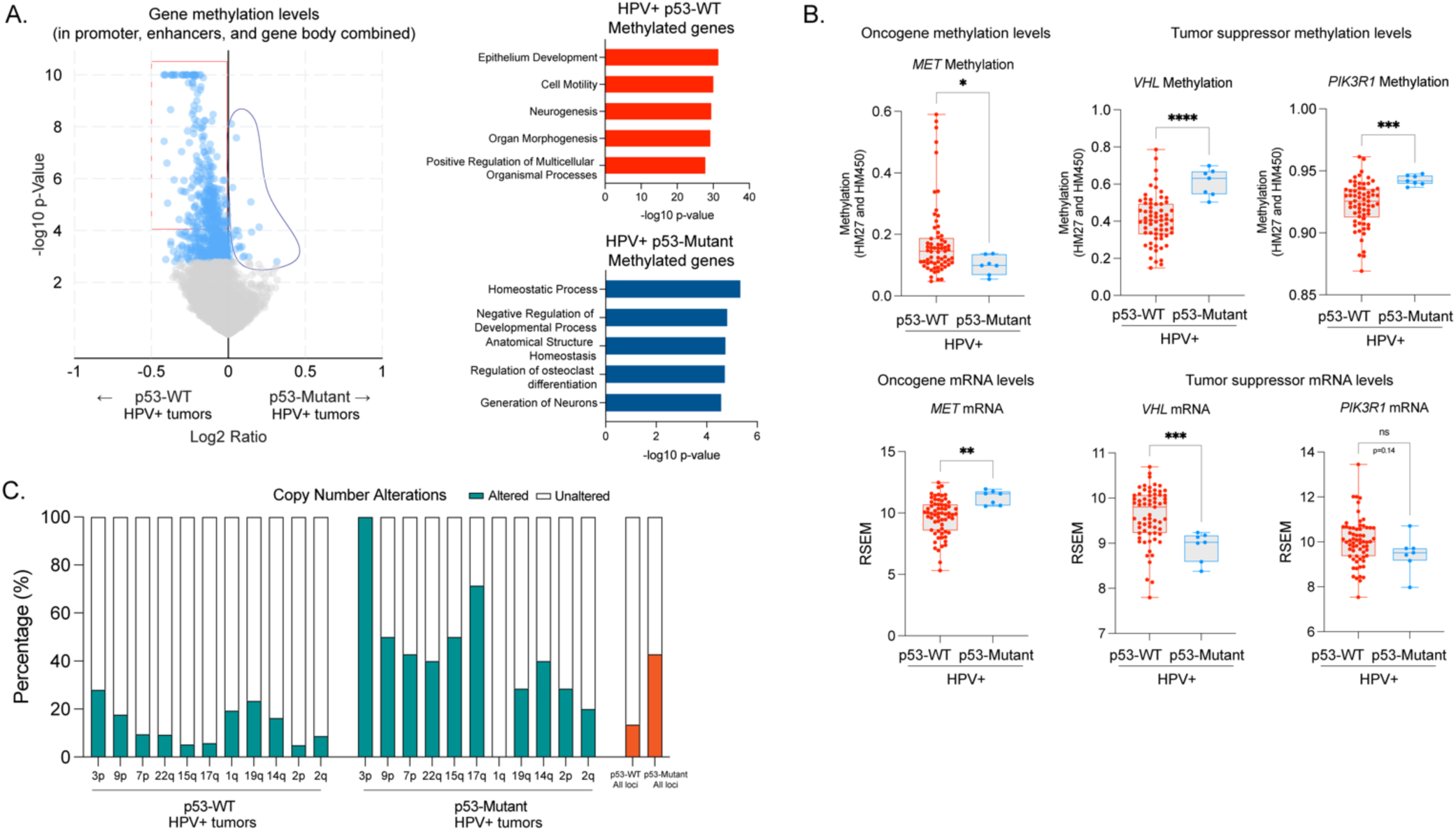
HPV+ p53-WT HNSCC tumors exhibit higher gene methylation levels and fewer chromosomal abnormalities compared to HPV+ p53-mutant tumors. **A.** Differential gene methylation is presented as log₂ fold change (log₂Ratio). In the volcano plot, significantly differentially methylated genes are highlighted in light blue. Genes with significantly higher methylation in *TP53*-WT tumors that were used for pathway enrichment analysis (500 genes) are outlined in red, while those with significantly higher methylation in *TP53*-mutant tumors used for pathway analysis are outlined in blue (23 genes). Right panels display pathway enrichment analysis of differentially methylated genes for each tumor type, based on the Gene Ontology Biological Process (GOBP) dataset. **B.** Normalized methylation levels (top row) and RNA-seq read counts (RSEM) (bottom row) for *MET, VHL,* and *PIK3R1* in HPV+ *TP53*-WT and HPV+ *TP53*-mutant HNSCC tumors. Statistical significance is indicated as follows: *p<0.05, **p<0.005, ***p<0.0005, ****p<0.00005, ns=not significant. **C.** Chromosomal alterations (gains and losses) across specified loci. The Y-axis represents the percentage of tumors with chromosomal alterations, while the X-axis denotes the specific genomic loci analyzed. Summary results (all loci) are shown to the right end of the plot (orange bars).

To determine whether differential methylation correlated with gene expression, we examined key oncogenes and tumor suppressors. The *MET* oncogene exhibited significantly higher methylation in *TP53*-WT tumors, which corresponded to lower *MET* mRNA expression in these tumors (Figure 5B). Conversely, *VHL* and *PIK3R1*, both tumor suppressors, showed significantly lower methylation levels and presented higher mRNA expression in *TP53*-WT compared to *TP53*-Mutant tumors (Figure 5B). These findings highlight the potential regulatory role of p53 in shaping the epigenetic landscape of HPV+ HNSCC tumors, influencing both oncogene suppression and tumor suppressor activation.

Lastly, given the role of p53 in maintaining genome integrity and our observation of significant upregulation of genes within the DNA repair pathway in *TP53*-WT HPV+ HNSCC tumors (Supplementary Figure 4), we investigated whether the incidence of chromosomal alterations is different in HPV+ *TP53*-WT compared to HPV+ *TP53*-mutant HNSCC tumors (Figure 5C). We identified significant differences in 11 genomic loci, with 10 of them exhibiting a higher frequency of alterations in *TP53*-mutant tumors. On average, fewer than 20% of *TP53*-WT tumors displayed chromosomal abnormalities (all loci), whereas over 40% of *TP53*-mutant tumors exhibited such alterations. These findings support a role for WT p53 in preserving genomic stability in HPV+ HNSCC, a key tumor suppressive mechanism.

### WT p53 suppresses PI3K-AKT signaling in HPV+ HNSCC

One of the p53 target genes identified earlier (Figures 3-4), *INPP5D*, encodes a phosphatase that converts PIP3 to PIP2, thereby inhibiting PI3K-AKT signaling. We found *INPP5D* to be significantly upregulated in HPV+ *TP53*-WT tumors (Figure 6A), suggesting a potential role for p53 in suppressing PI3K-AKT signaling.

**Figure 6.**
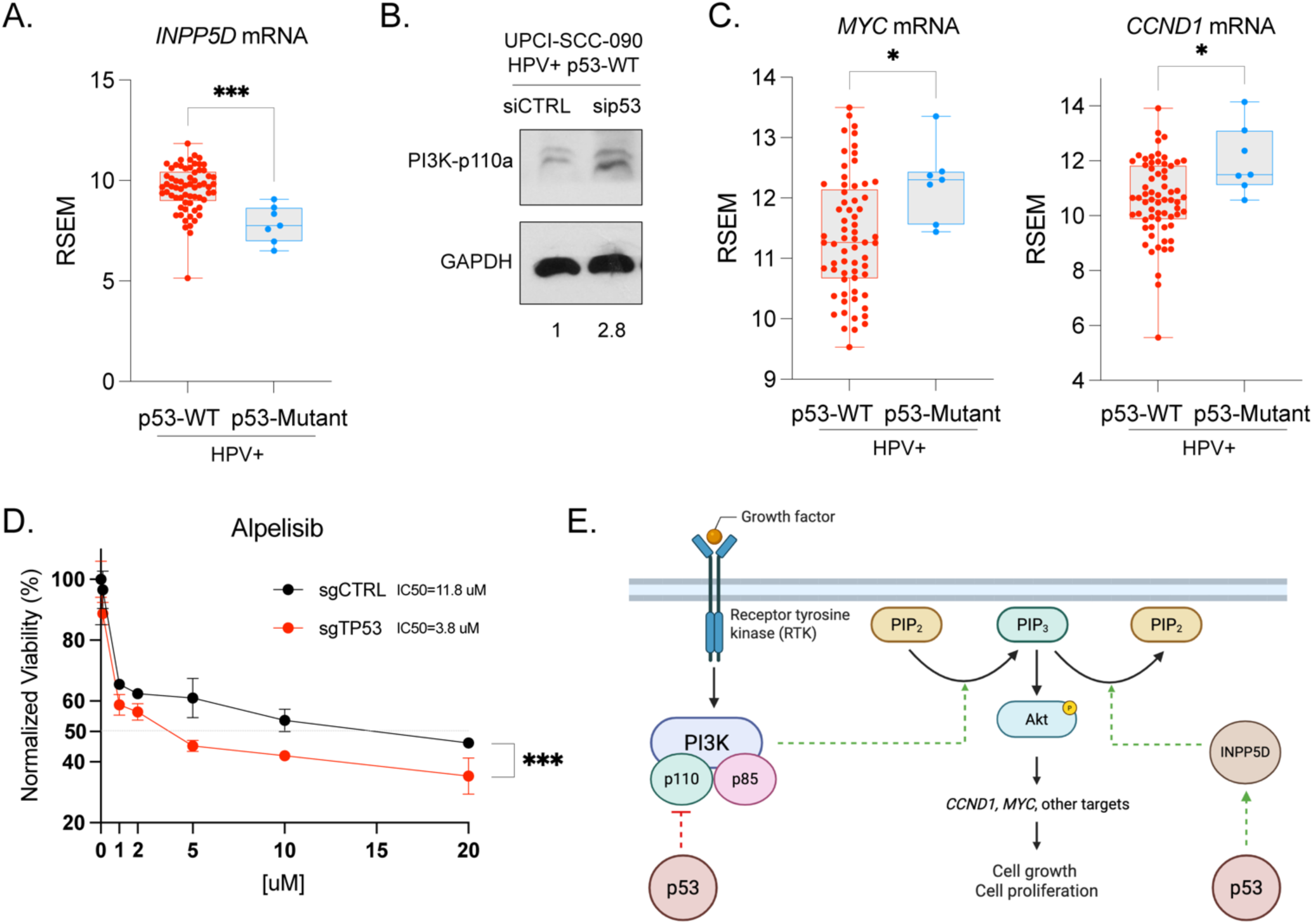
WT p53 suppresses the PI3K pathway in HPV+ HNSCC. **A.** Normalized RNA-seq read counts (RSEM) for *INPP5D* in HPV+ *TP53*-WT and HPV+ *TP53*-mutant HNSCC tumors. **B.** Immunoblot analysis of PI3K-p110α and GAPDH in UPCI-SCC-090 cells 48 hours post-transfection with siRNA targeting p53 (sip53) or a non-targeting control (siCTRL). Quantification of p110-α signal is shown relative to GAPDH and siCTRL. **C.** Normalized RNA-seq read counts (RSEM) for *MYC,* and *CCND1* in HPV+ *TP53*-WT and HPV+ *TP53*-mutant HNSCC tumors. **D.** Viability assay in UM-SCC-47-dCas9 cells with (sgCTRL) or without p53 (sgTP53) in the presence of increasing concentrations of Alpelisib (BYL719; 0-20 uM) **E.** Schematic representation of p53-mediated inhibition of the PI3K-AKT pathway in HPV+ HNSCC. *p<0.05, ***p<0.005.

Knockdown of WT p53 in the HPV+ HNSCC cell line UPCI-SCC-090 led to increased p110-α expression (PI3K catalytic subunit), indicative of PI3K pathway activation in the absence of WT p53 in HPV+ HNSCC (Figure 6B).

Further supporting this finding, we observed significantly increased *MYC* and *CCND1* mRNA levels - markers of AKT activation - in HPV+ *TP53*-mutant tumors compared to HPV+ *TP53*-WT tumors (Figure 6C).

Considering the historical failure of PI3K inhibitors in HPV+ HNSCC models, we investigated whether p53 ablation could enhance sensitivity to the FDA-approved PI3K inhibitor BYL719 (Alpelisib). Notably, UM-SCC-47 cells expressing dCas9 and treated with a sgRNA targeting p53 were more sensitive (IC50=3.8 uM) compared to cells treated with a non-targeting sgRNA (IC50=11.8uM) (Figure 5D).

Together, these findings suggest that WT p53 in HPV+ HNSCC suppresses PI3K signaling via *INPP5D* transactivation and p110-α inhibition (Figure 6E).

## Discussion

Patients with HPV+ HNSCC exhibit significantly better outcomes compared to patients with HPV-negative HNSCC [4]. A seminal discovery over 30 years ago revealed that HPV16 and HPV18-encoded E6 proteins promote p53 degradation [25]. Subsequent studies have shown that HPV may also impair p53 transcriptional activity [26]. These HPV-driven mechanisms of p53 inhibition parallel the physiological regulation of p53 by MDM2. Despite MDM2-mediated inhibition, p53 maintains homeostatic signaling critical for various cellular processes, including tumor suppression. Notably, while the majority of HPV+ HNSCC cases retain WT *TP53* alleles, 10% of cases harbor inactivating *TP53* mutations, suggesting a need to further dampen this pathway. Although it is well established that HPV suppresses p53 function, the extent of this suppression remains undefined. Here, we tested the hypothesis that HPV+ HNSCC retains residual p53-driven tumor suppressive functions.

To assess this, we first analyzed HPV+ HNSCC cases stratified by *TP53* status. Surprinsingly, we found that HPV+ tumors retaining WT *TP53* were associated with significantly better progression-free survival (median survival of ∼56 months) compared to both HPV+ *TP53*-mutant and HPV-negative tumors. The latter two groups showed no significant differences, with comparable median survival times of 15 and 19 months, respectively. This finding suggests that the long-recognized survival advantage of HPV+ over HPV-negative HNSCC is primarily driven by the presence of a WT *TP53* allele.

To investigate potential mechanisms underlying this finding, we further inhibited p53 expression in HPV+ *TP53*-WT HNSCC cell lines via siRNA or CRISPRi and conducted functional assays. Consistent with our hypothesis, p53 knockdown enhanced cell proliferation, migration, and invasion, indicating that residual p53 activity suppresses tumor-promoting phenotypes even in the presence of HPV. Transcriptomic analyses further revealed that p53 continues to regulate gene expression, activating tumor suppressor genes (e.g. *INPP5D*) while repressing tumor-promoting genes (e.g. *RRM2*). Notably, beyond its well-established role in cell cycle control, p53-regulated genes were enriched in pathways related to metabolism and inflammation, highlighting its broader regulatory influence on cellular functions.

E6 knockdown effectively elevated expression of the p53 targets *CDKN1A* and *INPP5D*, suggesting that this is a p53-dependent phenotype. Using a pair of HPV-negative isogenic cell lines, we further demonstrated that many of the genes regulated by p53 in HPV+ cells are also regulated by p53 in the absence of HPV. These findings collectively support a model in which HPV suppresses but does not fully abolish p53 function in HNSCC.

To extend these findings to patient tumors, we analyzed molecular features of HPV+ HNSCC cases stratified by *TP53* status. We examined the expression of p53-regulated genes identified in our cell line models within HPV+ HNSCC tumors. Among 15 genes tested, only *INPP5D* (also known as SHIP1), an understudied direct p53 target [27, 28] and tumor suppressor, was found to be significantly upregulated in *TP53*-WT compared to *TP53*-mutant HPV+ tumors. Given that INPP5D negatively regulates PI3K-AKT signaling, we assessed pathway activity and observed increased expression of *MYC* and *CCND1* in *TP53-*mutant tumors, consistent with enhanced PI3K-AKT activity. Supporting this observation, the PI3K subunit p110-α was upregulated upon WT p53 knockdown in HPV+ HNSCC cells, further implicating p53 in restraining PI3K signaling.

These findings led us to hypothesize that WT p53 functions as an endogenous “brake” on PI3K pathway activation, and that loss of p53 may sensitize cells to pharmacologic PI3K inhibition.

Consistent with this, p53 ablation in HPV+ HNSCC cells significantly increased sensitivity to Alpelisib, reducing the IC50 by ∼3 fold (IC50=11.8 uM in sgCTRL vs 3.8 uM in sgTP53).

These findings have important implications as HPV+ HNSCC is generally resistant to PI3K-targeting therapies [29]. Our data provide a potential mechanistic explanation: residual WT p53 activity may partially suppress PI3K signaling, reducing cellular dependence on this pathway and thereby limiting the efficacy of PI3K inhibitors. In contrast, *TP53*-mutant HPV+ tumors, which exhibit upregulation of the PI3K-AKT pathway (potentially due to complete loss of p53 function), may remain sensitive to PI3K inhibition. This raises the hypothesis that stratifying HPV+ HNSCC patients by *TP53* status could improve treatment outcomes by identifying patients who are more likely to benefit from PI3K-targeted therapies. Future studies should explore whether *TP53*-mutant HPV+ tumors display increased sensitivity to PI3K inhibitors in clinical settings, potentially leading to more personalized treatment strategies.

Further investigation of the molecular features of HPV+ HNSCC tumors based on *TP53* status revealed a higher degree of gene methylation in *TP53*-WT cases compared to *TP53*-mutant tumors. Notably, many of the hypermethylated (silenced) genes in *TP53*-WT tumors were tumor-promoting, including the oncogene *MET*, which correlated with lower *MET* mRNA expression in these tumors. Beyond differences in methylation and gene expression, *TP53*-WT tumors also exhibited fewer chromosomal abnormalities.

Collectively, our findings support a new paradigm of residual p53 activity in HPV+ HNSCC, with significant clinical implications. These findings highlight the existence of molecularly distinct subtypes within HPV+ HNSCC defined by *TP53* status. While the majority of HPV+ HNSCC cases are associated with favorable clinical outcomes, a subset displays aggressive behavior resembling that of HPV-negative tumors. Based on our findings, we propose that this aggressive HPV+ subset is characterized by the presence of *TP53* mutations, which may override the tumor-suppressive effects of residual WT p53 activity.

Consequently, therapeutic strategies should not be uniform for all HPV+ HNSCC cases but rather tailored based on *TP53* status. For instance, MET inhibitors such as cabozantinib may be more effective in *TP53*-mutant HPV+ HNSCC, where MET is more highly expressed, whereas *TP53*-WT HPV+ tumors may derive less benefit due to epigenetic silencing of *MET*. Additionally, whether residual p53 signaling contributes to the observed resistance of HPV+ HNSCC to Cetuximab [30], a widely used EGFR-targeting therapy, remains an open question. Similarly, the role of WT p53 in mediating responses to radiotherapy in HPV+ HNSCC warrants further investigation. Our data also suggest that WT p53 may modulate immune responses in HPV+ HNSCC (Figure 3, Supplementary Figure 4), highlighting the potential for stratifying immune checkpoint inhibitor (ICI) therapy by *TP53* status. Further molecular studies are needed to mechanistically define the role of p53 in these therapeutic responses, an important consideration for future treatment strategies.

Lastly, this study offers a new perspective on HPV+ *TP53*-WT HNSCC models, positioning them as valuable systems to explore HNSCC-specific functions of p53. Here, we identified *INPP5D* as a key p53 target that is a suppressor of PI3K signaling, revealing a novel mechanism of tumor suppression. However, a deeper mechanistic understanding of how p53 controls genomic stability, DNA methylation patterns, and other tumor-relevant processes in HNSCC may uncover additional targetable vulnerabilities and expand therapeutic opportunities.

### Conclusion

Together, our findings reveal that p53 retains functional relevance in HPV+ HNSCC and may significantly influence both tumor behavior and therapeutic response. Understanding the molecular distinctions between *TP53*-WT and *TP53*-mutant HPV+ HNSCC could enhance patient stratification and guide the development of more effective, precision-based therapies. Future studies should aim to elucidate how residual p53 activity modulates key oncogenic pathways and whether targeting these pathways can improve outcomes in *TP53*-mutant HNSCC. Importantly, HPV+ *TP53*-WT HNSCC models represent a powerful platform for uncovering disease-specific, p53-driven tumor-suppressive mechanisms - offering potential for pharmacologic reactivation of these pathways in *TP53*-mutant HNSCC to improve patient outcomes.

## Materials and Methods

### Cell lines and cell culture

Human cell lines UM-SCC-47, UPCI-SCC-090 and UD-SCC-2 were selected from a panel of HPV+ HNSCC cell lines that have no endogenous *TP53* mutation as confirmed by the Broad Institute Cancer Cell Line Encyclopedia. UM-SCC-47 cells engineered to express dCas9 are described elsewhere [31]. These cells were maintained in Dulbecco’s Modified Eagle’s medium (Corning) supplemented with 10% fetal bovine serum, and 1% penicillin/streptomycin. Cells were maintained in a humidified incubator at 37°C and 5% carbon dioxide. Each cell line was thawed and passaged approximately 3-4 times before necessary experimentation. Genotype authentication for human cell lines was performed by LabCorp using the STR method (last authenticated on 08/06/2024). Isogenic human HPV-negative PCI-13 cell lines (p53WT and p53G245D) were obtained from the laboratory of Dr. Jeffrey N. Myers at The University of Texas MD Anderson Cancer Center and maintained as described previously [24].

### Proliferation, migration, invasion and drug susceptibility assays

*Proliferation:* cells were seeded at approximately 10,000 cells per well in 48 well plates. 24 hours after seeding, cells were transfected with siRNA or sgRNA constructed to target p53 (Horizon Discovery, L- 003329-00-0005 or CF-003329-01-0005, respectively) or non-targeting control (Horizon Discovery, D-001810-10-20 or U-009550-10-05) using Lipofectamine RNAiMAX (ThermoFisher, 13778075). Cells were incubated for 48 hours (UM-SCC-47 and UD-SCC-2), 72 hours (UPCI-SCC-090), or 120 hours (UM-SCC-47-dCas9) at 5% CO2 and 37 degrees. Crystal violet assay was performed as directed by manufacturer (Abcam Kit, ab232855).

*Migration and invasion:* cells were seeded at 300,000 cells per well in 6-well plates. 24 hours after seeding, cells were transfected with siRNA constructed to target p53 (Horizon Discovery, L-003329-00-0005) or non-targeting control (Horizon Discovery, D-001810-10-20) using Lipofectamine RNAiMAX (ThermoFisher, 13778075). 24 hours after transfection, cells were harvested, counted, and re-seeded in FBS-free media in either 24-well Migration plates (polycarbonate membrane) or 24-well Invasion Chambers (basement membrane) (50,000 cells/well) as directed by the manufacturer (CellBioLabs INC, CBA-100-C). We used 10% FBS-supplemented growth media as attractant. Twenty hours later, we measured cell migration and invasion using the provided cell viability staining solution (CellBioLabs INC, CBA-100-C). Images were taken using a light microscope following staining.

*Drug susceptibility:* UM-SCC-47-dCas9 cells were seeded at approximately 300,000 cells per well in 6-well plates. 24 hours after seeding, cells were transfected with sgRNA constructed to target p53 (Horizon Discovery, CF-003329-01-0005) or non-targeting control (Horizon Discovery, U-009550-10-05) using Lipofectamine RNAiMAX (ThermoFisher, 13778075). After another 24 hours, cells were trypsinized, counted, and re-seeded at 10,000 cells per well in 48-well plates. The next day, growth media was replaced by media containing varying concentrations of the PI3K inhibitor Alpelisib (BYL719, Sigma). Cells were incubated for 72 hours in the presence/absence of drug at 5% CO2 and 37 degrees. Crystal violet assay was performed at end point as directed by manufacturer (Abcam Kit, ab232855).

### Immunoblotting

Cells were seeded at 300,000 cells per well in a 6-well plate. 24 hours after seeding, cells were transfected with siRNA constructed to target p53 (Horizon Discovery, L-003329-00-0005), E6 (Santa Cruz, sc-156008) or non-targeting control (Horizon Discovery, D-001810-10-20) using Lipofectamine RNAiMAX (ThermoFisher, 13778075). Cells were harvested at 24, 30, 48, or 72 hours post-transfection using cell lysis buffer (Cell Signaling Technology, Cat#9803 with cOmplete Protease Inhibitor Cocktail (Roche, Cat#11836170001). Protein concentration was determined using a Bradford Assay (Bio-Rad, Cat#5000006). Proteins were standardized to 100 ug/uL, diluted with cell lysis buffer. 30 ug of protein was electrophoresed on a 10% SDS-PAGE gel and transferred to a PVDF membrane (Bio-Rad, Cat#1620177) at 25V for 1 hour. Membranes were blocked in 5% nonfat dried milk (Apex Bioresearch, Cat#20-241) reconstituted in TBST (Teknova, Cat#T1680) for 15 minutes. Membranes were incubated in a 1:1000 concentration of primary antibodies overnight at 4 degrees. Primary antibodies: GAPDH (D16H11) XP® Rabbit mAb (Cell Signaling Technology, Cat # 5174S), p53 (DO1) (Santa Cruz Biotechnology, Cat# sc-126), p21 Wafl/cipl (12D1) (Cell Signaling Technology, Cat# 2947S), CDK1/CDK2 (Santa Cruz Biotechnology, Cat# sc-53219), PI3K p110-alpha (Cell Signaling Technology, Cat# 4255). Membranes were incubated in 1:3000 concentration of secondary HRP-labeled antibody in 2.5% nonfat dried milk for 2 hours at room temperature. Membranes were developed using Luminol Reagent (Santa Cruz Biotechnology, Cat# sc-2048) and imaged using the Bio-Rad ChemiDoc Imaging system. ImageJ (version 1.54; National Institutes of Health, Bethesda, MD) was used for densitometry analysis.

### Quantitative RT-PCR

Cells were seeded at 300,000 cells per well in a 6-well plate. 24 hours after seeding, cells were transfected with siRNA constructed to target p53 (Horizon Discovery, L-003329-00-0005), E6 (Santa Cruz, sc-156008), or non-targeting control (Horizon Discovery, D-001810-10-20) using Lipofectamine RNAiMAX (ThermoFisher, 13778075). Cells were harvested at 8, 12, 24, 30, or 48 hours post-transfection, and RNA was isolated using the Direct-zol RNA MiniPrep with TRIReagent kit (Zymo Research, Cat# 2051). RNA concentrations were measured using a NanoDrop spectrophotometer (Thermo Fisher). 1 ug of RNA was used to generate cDNA using iScript Reverse Transcription Supermix (Bio-Rad, Cat# 1708841). An Eppendorf Mastercycler Nexus Thermal Cycler (Millipore Sigma, Cat# EP6334000026) was used to generate cDNA. Tripplicate or quadruplicate biological samples were prepared in technical triplicates in 96-well RT-qPCR plates (Bio-Rad, Cat#HSP9631). Primers were designed and purchased from Integrated Data Technologies. Primers are as follows: human *CDKN1A (p21)* (F: 5’-AGGTGGACCTGGAGACTCTCAG-3’; R: 5’-TCCTCTTGGAGAAGATCAGCCG-3’), human *INPP5D* (F: 5’-TGTGACCGAGTCCTCTGGAAGT-3’; R: 5’-GCCTCAAATGTGGCAAAGACAGG-3’), human TM7SF3 (F: 5’-CCAGCAAACCTAGGCTATGCGA-3’; R: 5’-GCAACATCTCCTCAGTGAGGTC-3’) and human *ACTINB* (F: 5’-CACCATTGGCAATGAGCGGTTC-3’; R: 5’-AGGTCTTTGCGGATGTCCACGT-3’). *ACTINB* mRNA expression levels were used as the internal control. Expression levels were calculated using the double delta cq method. Graphs were constructed using PRISM 10 Graphpad software.

### RNA-sequencing and analysis

UM-SCC-47 and UPCI-SCC-90 cells were seeded at 300,000 cells per well in 6-well plates. After 24 hours, cells were transfected with either p53-targeting siRNA (Horizon Discovery, L-003329-00-0005) or a non-targeting control (Horizon Discovery, D-001810-10-20) using Lipofectamine RNAiMAX (Thermo Fisher, 13778075). Cells were harvested 30 hours post-transfection, and RNA was extracted using the Direct-zol RNA MiniPrep with TRI Reagent kit (Zymo Research, Cat#2051). RNA concentrations were measured with a NanoDrop spectrophotometer (Thermo Fisher) and two biological replicates per condition were sent to Novogene for sequencing (Illumina PE150, 20M paired reads). Bioinformatic analysis was performed by Novogene using the DESeq2 software package for differential gene expression analysis. Genes with p-values < 0.01 were considered significant, regardless of q-values. Volcano plots were generated in RStudio.

FASTQ RNA-seq data files from isogenic PCI-13 p53WT and PCI-13 p53G245D cells were provided by Dr. Jeffrey N. Myers (The University of Texas MD Anderson Cancer Center) under a Material Transfer Agreement. Reads were aligned using the nf-core/RNAseq pipeline in paired-end mode and normalized read counts were generated and analyzed in R using limma, edgeR, and voom.

### Human HNSCC tumor data

All data related to human HNSCC tumors including survival, gene expression, gene methylation, and chromosomal abnormalities, was obtained from cBioPortal [2] using the TCGA PanCancer Atlas dataset.

### Pathway Enrichment Analysis

We used the Gene Set Enrichment Analysis (GSEA) online tool [32] to identify significantly enriched pathways in differentially expressed or differentially methylated genes. Enrichment analysis was performed using MSigDB GOBP, Reactome, and Wikipathways gene sets, with statistical significance determined by FDR q-value < 0.05.

### Statistical analysis

Most data were analyzed using GraphPad Prism version 10, unless otherwise specified in the respective methods section. For cell line experiments, comparisons between two groups were performed using unpaired two-tailed Student’s t-tests. Human tumor data comparisons were conducted using the Mann-Whitney U test (Wilcoxon rank-sum test). Survival analysis was performed using the log-rank (Mantel-Cox) test. IC50 values were determined by non-linear regression using a log(inhibitor) vs. normalized response curve (variable slope model). Comparisons of IC50 values between conditions were based on curve fitting and inspection of confidence intervals. A p-value < 0.05 was considered statistically significant.

## Author Contributions

J.G-A. conceptualized the study and performed experimentation, data analysis, data interpretation, and writing of original manuscript. H.L. contributed to experimentation. L.C.W. contributed to experimentation and writing of original manuscript. A.A.B. conducted bioinformatic analysis of RNA-seq files. J.N.M. provided cell lines, RNA-sequencing data, and edited original manuscript. P.H., D.E.J and J.R.G, supervised, secured funding, and edited original manuscript.

## Acknowledgements

This work was supported by grants R35 CA231998 and a Cancer Cell Map Initative (CCMI) grant U54 CA209891. J.G-A. is supported by a University of California President’s Postdoctoral Fellowship and a Pew Charitable Trusts Biomedical Sciences Fellowship.

## Supplemental Figures

**Supplemental Figure 1.**
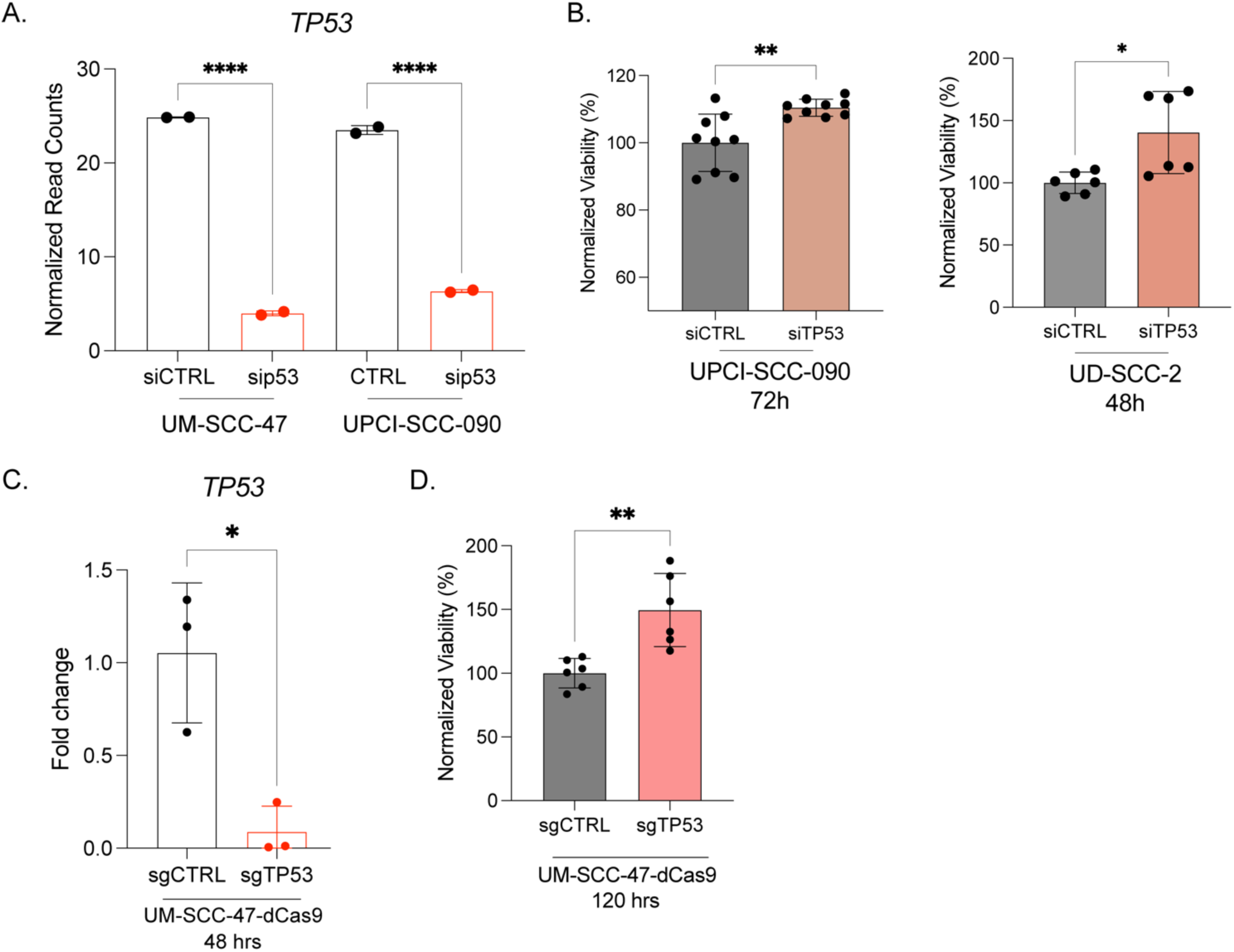
*TP53* mRNA levels and cell viability assessment following p53 knockdown. **A.** *TP53* mRNA levels in UM-SCC-47 and UPCI-SCC-090 cells transfected with either non-targeting siRNA (siCTRL) or siRNA targeting p53 (sip53). RNA was collected 30 hours post-transfection, and RNA sequencing analysis was performed. Data are presented as normalized read counts. **B.** Viability assays following p53 knockdown (72 hours for UPCI-SCC-090; 48 hours for UD-SCC-2). Data are normalized to siCTRL and plotted as mean ± SEM. **C.** *TP53* mRNA levels (RT-qPCR) in UM-SCC-47-dCas9 48 hours post-transfection with sgRNA targeting p53 (sgTP53) or non-targeting control (sgCTRL). **D.** Viability assays following p53 knockdown (CRISPRi) at 120 hours post-transfection with sgRNA targeting p53 (sgTP53) or non-targeting control (sgCTRL). Statistical significance: *p < 0.05, **p < 0.005, ****p < 0.00005.

**Supplemental Figure 2.**
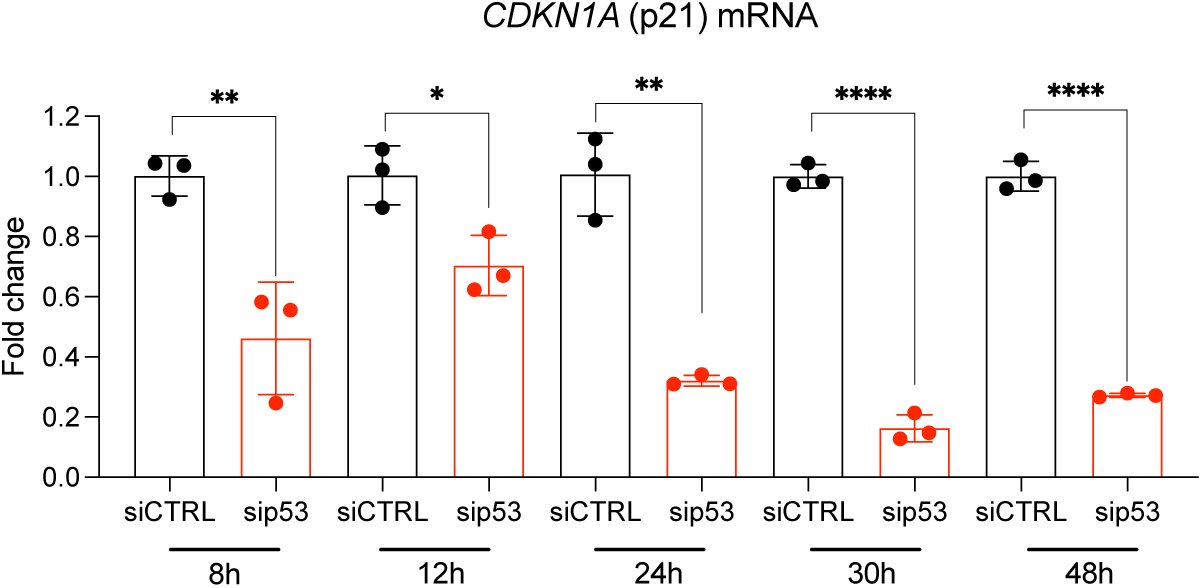
*CDKN1A* mRNA levels are maximally decreased at 30 hours following p53 knockdown. UM-SCC-47 cells were transfected with either non-targeting siRNA (siCTRL) or siRNA targeting p53 (sip53). RNA was collected at 8, 12, 24, 30, and 48 hours post-transfection. qPCR analysis was performed to quantify *CDKN1A* mRNA levels, normalized to *ACTB* and expressed relative to siCTRL. Data are plotted as mean ± SEM. Statistical significance: *p < 0.05, **p < 0.005, ****p < 0.00005.

**Supplemental Figure 3.**
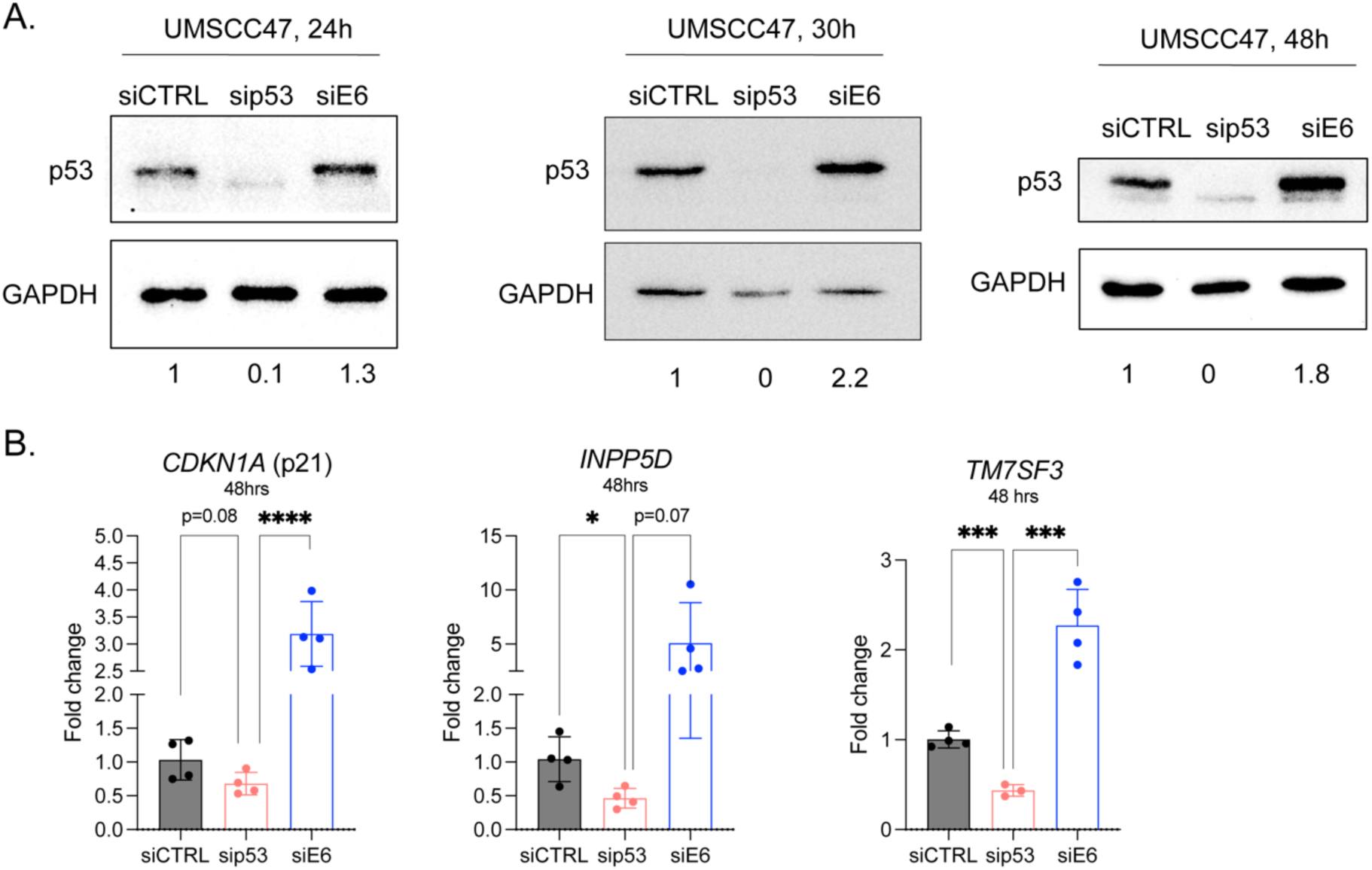
p53 protein levels and p53 target gene expression increase following E6 knockdown. **A.** UM-SCC-47 cells were transfected with either non-targeting siRNA (siCTRL), siRNA targeting p53 (sip53), or siRNA targeting E6 (siE6). Cell pellets were collected at 24, 30, and 48 hours post-transfection. Immunoblots display p53 and GAPDH protein levels, with quantifications below showing relative p53 protein levels, normalized to GAPDH (loading control) and expressed relative to siCTRL conditions. The upper (darker) band in each p53 blot represents the p53-specific signal. **B.** mRNA expression levels of p53 target genes *CDKN1A*, *INPP5D, TM7SF3* decrease upon p53 knockdown and are restored (and elevated) following E6 knockdown in HPV+ HNSCC (UM-SCC-47) cells 48 hours post-transfection.

**Supplemental Figure 4.**
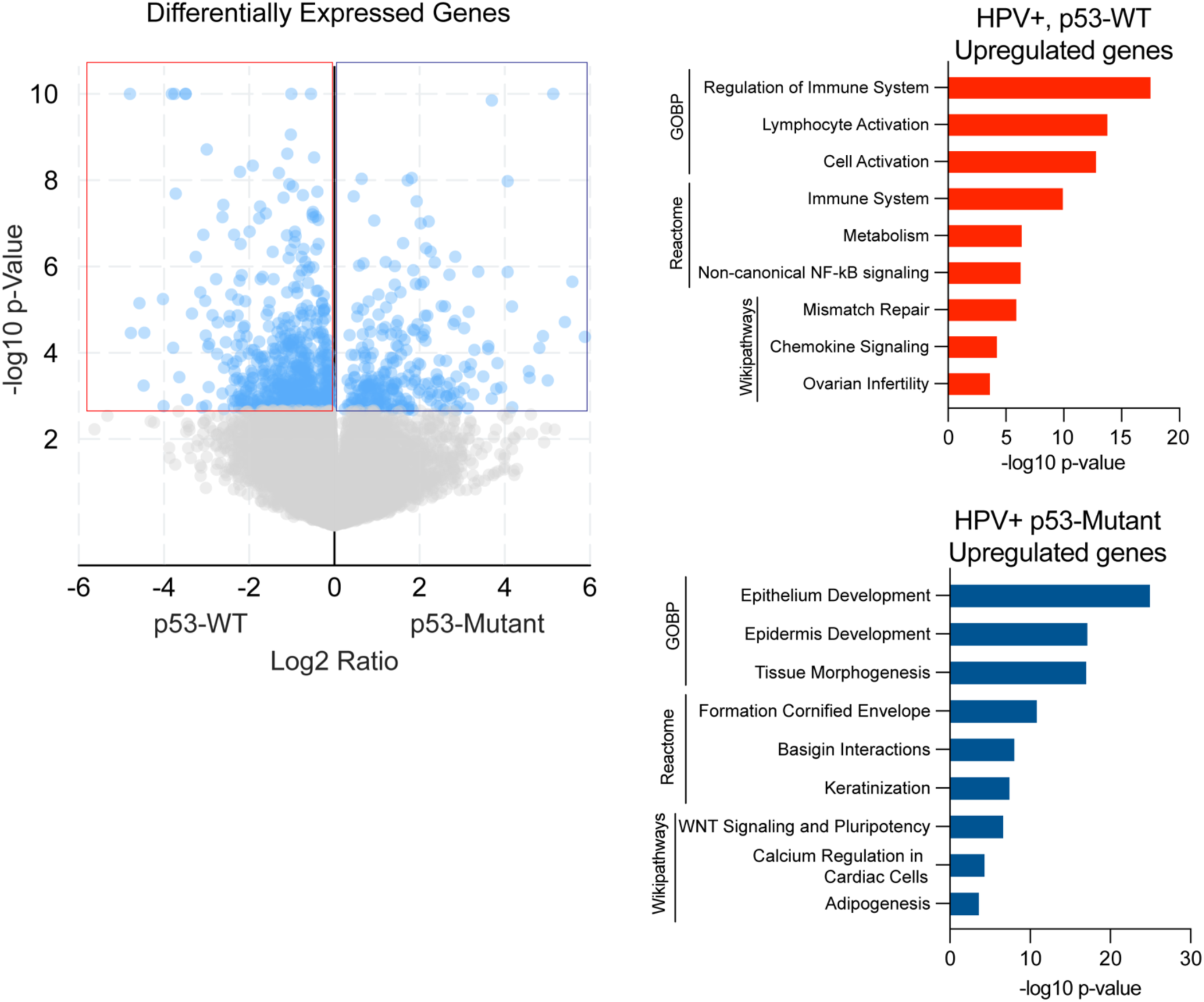
Global transcriptomic differences between HPV+ *TP53*-WT and HPV+ *TP53*-mutant HNSCC tumors. Differential gene expression is presented as log₂ fold change (log₂Ratio). Significantly differentially expressed genes (DEGs) are highlighted in light blue in volcano plot. Red outline indicates genes significantly upregulated in *TP53*-WT tumors, while the blue outlined box represents genes significantly upregulated in *TP53*-mutant tumors. Right panels show pathway enrichment analysis of DEGs for each tumor type, with the top three significant pathways from three datasets plotted (GOBP, Reactome, Wikipathways). Data extracted from TCGA (cBioPortal). GOBP=Gene Ontology Biological Process dataset.

